# Longitudinal changes in brain activation underlying reading fluency

**DOI:** 10.1101/2021.07.09.451857

**Authors:** Ola Ozernov-Palchik, Dana Sury, Ted K. Turesky, Xi Yu, Nadine Gaab

**Affiliations:** McGovern Institute for Brain Research, Massachusetts Institute of Technology, Cambridge, Massachusetts, USA; Harvard Graduate School of Education, Harvard University, Cambridge, Massachusetts, USA; Laboratories of Cognitive Neuroscience, Boston Children’s Hospital; Harvard Medical School; State Key Laboratory of Cognitive Neuroscience and Learning, Beijing Normal University

## Abstract

Reading fluency – the speed and accuracy of reading connected text – is foundational to educational success. The current longitudinal study investigates the neural correlates of fluency development using a connected-text paradigm with an individualized presentation rate. Twenty-six children completed a functional MRI task in 1st/2nd grade (*time 1*) and again 1-2 years later (*time 2*). There was a longitudinal increase in activation in the ventral occipito-temporal (vOT) cortex from *time 1* to *time 2*. This increase was also associated with improvements in reading fluency skills and modulated by individual speed demands. These findings highlight the reciprocal relationship of the vOT region with reading proficiency and its importance for supporting the developmental transition to fluent reading. These results have implications for developing effective interventions to target increased automaticity in reading.

## Introduction

Reading fluency is the foundation for proficient reading and is critical to educational success (NRP, 2000). The term *fluency* refers to the speed and accuracy of decoding connected text (Chard, Vaughn, & Brenda-Jean Tyler, 2002). Despite extensive research into the brain basis of reading, the topic of fluency development has been largely overlooked in the neuroimaging literature. Insights into the neural processes underlying fluency development are important for understanding fluency deficits in children with reading difficulties and for the development of effective interventions targeting these deficits. The current longitudinal study investigates the neural correlates of fluency using a connected-text paradigm during a period of time in which children transition from non-fluent to fluent reading.

The goal of successful reading acquisition is to read an unfamiliar text fluently, with great automaticity, and comprehend it. In typical reading development in English-speakers, children acquire fluency in grades 2 and 3 (roughly 8 to 9 years old; Chall, 1996; Indrisano & Chall, 1995). The development of fluency has been conceptualized as the outcome of achieving proficiency in the lower-level component skills of reading (Kame’enui, Simmons, Good, & Harn, 2001, Wolf & Katzir-Cohen, 2001). More specifically, fluency is achieved when processing at the phonological, orthographic, semantic, and morphological levels -- and critically, among these levels -- becomes automatic. Automaticity has been defined as processing without expending attention or effort (Ehri, 2005, p. 151). Automaticity arises as a result of robust associations being formed between written words and their linguistic representations (i.e., phonological and semantic) through learning and practice (Ehri, 2005; 2011; Hudson et al., 2008). Once these associations are established, word identification of familiar words becomes primarily a memory retrieval process that proceeds quickly and without reader’s conscious control, resulting in fluent reading of connected text. This allows for processing words in a fashion that support connecting words together into meaningful strings and for allocating cognitive resources to support processes related to comprehension of text (Perfetti, 1985).

Fluency serves as the foundation for the next stage in reading development -- reading to learn -- that occurs in later grades (Chall, 1996). When word recognition is not efficient, cognitive resources that are needed to support text integration and comprehension are instead deployed to support word identification (Crain & Shankweiler, 1990; Ozernov-Palchik et al., 2020; Wolf & Katzir-Cohen 2001). Indeed, there is evidence that fluency makes a unique contribution to reading comprehension beyond accuracy (Cutting et al., 2009; Joshi & Aaron, 2000; Silverman et al., 2013; Tilstra et al., 2009) and has important implications for children with reading difficulties, as a fluency deficit may describe some of the most impaired readers, particularly in older grades (Wolf & Bowers, 1999). Thus, fluency is a critical prerequisite for reading comprehension, but the neurocognitive processes underlying the development of fluency remain relatively unknown.

Neuroimaging studies of reading development have demonstrated that foundational reading skills such as mapping phonemes (i.e., speech sounds) to their orthographic representations (i.e., letters) are associated with the structure and function of the temporoparietal brain regions. A shift from early-reading in English (5-6 years) to emergent reading (7-8 years) and subsequently increasingly fluent reading (8-9 years) has been associated with increased development and recruitment of the occipito-temporal brain regions (Pugh et al., 2001; review by Chyl et al., 2021). The increased specialization of the ventral occipito-temporal cortex (vOT) for print has emerged as an important milestone for the development of word reading (Dehaene et al., 2015). In particular, increased response of the vOT region to words has been associated with better reading proficiency (Ben-Shachar et al., 2011 Brem et al., 2020; Kubota et al., 2019; Maurer et al., 2011; Olulade et al. 2013; Parviainen et al., 2006), and has been shown longitudinally in response to reading instruction and intervention (Brem et al., 2010; Fraga-González et al., 2015; Rezaie et al., 2011; Shaywitz et al., 2004) and in older as compared to younger readers (Ben Shachar et al., 2011; Smith, Booth, McNorgan, 2018).

As a result of its advantageous structural connections to the phonological, semantic, and memory systems in the brain, the vOT region becomes specialized for automatic word recognition with increased reading experience (Centanni et al., 2019; Dehaene et al., 2010, 2015; Dehaene and Cohen, 2007; Saygin et al., 2016; Stevens et al., 2017; Wang et al., 2020). For example, the connectivity of the vOT in pre-readers, but not its responsiveness to print, has been shown to predict the functional specificity of the region for words three years later (Saygin et al., 2016). In earlier stages of reading development, the vOT emerges as a hub linking visual letter patterns with first phonological and then semantic representations; with increased reading expertise, vOT assists in linking orthographic patterns directly with semantic representations. In fluent readers, this region is thought to process words in a similar way that other proximal regions in the left and right hemispheres process objects such as faces, identifying them wholistically and without exerting conscious effort (Dehaene and Cohen, 2007; Mei et al., 2010).

Despite the overall understanding of the development of the reading brain circuitry and the important role of vOT in automatic word recognition, it remains unknown how reading fluency develops in the brain. Studies investigating the brain correlates of reading have primarily focused on single-word or letter identification for their functional tasks (Aboud et al., 2018; Ben Shachar et al., 2011; Brem et al., 2010; Eden et al., 2004; Olulade et al. 2013; Shaywitz et al., 2004), or on reading comprehension without attending to reading fluency. Integrating across words while reading connected text, however, is an important feature of fluency during naturalistic reading. Several studies compared individuals with reading fluency deficits to typical readers using sentence-level stimuli and observed differences in activation in left temporoparietal (Meyler et al. 2007; Rimrodt et al. 2009; Schulz et al. 2009), occipito-temporal, and inferior frontal gyrus areas (Kronbichler et al. 2006). These studies, however, focused on sentence comprehension and were not longitudinal.

Longitudinal designs allow investigators to characterize the neural changes associated with a particular cognitive function in the same individuals. Although a limited number of longitudinal studies have used sentence tasks (Nugiel et al., 2019; Roe et al., 2018), these studies focused on measures of comprehension but not fluency and investigated brain differences in relation to intervention response, rather than to business-as-usual development and schooling. Furthermore, these studies held the speed of word processing constant, not accounting for individual differences in the rate of word processing, an important indicator of fluency (Chard, Vaughn, & Tyler, 2002). Therefore, no previous neuroimaging studies have used naturalistic sentence-level stimuli and manipulated individual reading speed to longitudinally investigate the neural substrates of fluency development.

A more ecologically-valid approach to neuroimaging of fluency was implemented in several previous studies that measured differences in patterns of activation when reading speed is manipulated within the same individuals and sentence-level stimuli are used (Benjamin & Gaab, 2012; Christodoulou et al., 2014; Kujala et al., 2007; Langer et al., 2015, 2019). For example, Langer et al. (2015; 2019) presented sentences at constrained, comfortable, and accelerated speeds determined based on individual reading speed to 8-12-year-old children with and without a reading disability. Both groups of children showed an increased response in bilateral vOT with increased fluency demands. Using the same task, another study in adult participants also reported increased activity in the vOT regions with higher speed demands (Benjamin & Gaab, 2012). A key finding from these studies is increased activation in the vOT cortex with increased reading speed; however, the developmental significance and timeline of these findings for emerging fluency remains undetermined.

The current study examined longitudinal changes in brain activation associated with fluent reading during the period in which children typically transition from early to fluent reading. All children underwent functional MRI while performing a reading fluency task (Benjamin & Gaab, 2011; Langer et al., 2013, 2019) in which the speed of text presentation was manipulated at both time points. A critical advantage of this approach for developmental research is controlling for task demands across reading proficiency levels. If text were presented at the same speed to all participants, slower readers (in this case younger readers) may be presented with a more challenging task than faster readers. This may result in increased recruitment of multi-demand domain-general brain regions, rather than regions that support reading fluency, the focus of this study. Therefore, comfortable reading speed was determined for each child prior to the scan at both time points, and this speed was used for the in-scanner task manipulation of two speeds of presentation: comfortable and accelerated.

Based on previous findings of increased engagement of vOT with reading proficiency and with increased speed demands, we hypothesized that we would (1) observe increased engagement of the vOT areas in older children as compared to younger children when comparing comfortable reading speeds; (2) increased engagement of these regions with increased reading speed demands in both age groups; and (3) an association between the increased activation in the vOT regions and improvement in reading fluency performance across the two time points.

## Methods

### Participants

Children (*N* = 26) were retrospectively selected from the Boston Longitudinal Dyslexia study (BOLD) aimed to study the neural trajectory underlying typical and atypical reading development in children with and without a family history of developmental dyslexia (e.g., Powers et al., 2016; Raschle et al., 2011, 2012; Wang et al., 2017; Yu et al., 2020). Only participants whose fluency neuroimaging task and behavioral data were successfully collected at two time points within a time gap of 1-2 years were included in the current study (*N* = 31). Five children who performed the in-scanner fluency task with less than 70% accuracy were excluded from analyses, resulting in a sample of typical developing children. As a result, twenty-six children (15 male) were included the final sample for the current study. The mean age was 8.25 years (SD = 9 months; children were in first or second grade) for the first time point and 9.5 years (SD = 14 months; children were in third or fourth grade) for the second time point, with a mean of 14±8 months between the two time points. All children were right-handed, native English speakers with no history of neurological symptoms, head injuries, visual problems, or hearing loss. The study was approved by the Institutional Review Board at Boston Children’s Hospital. Written informed consent was obtained from each participant’s accompanying parent, and verbal consent was obtained from each participant. Parental education information is summarized in **Supplemental Table 1**.

### Psychometric measurements

All children were examined using a comprehensive battery assessing language, pre-reading and reading skills. To avoid redundancy and reduce the number of comparisons, group characterization for the two time points focused on assessments that tested specific reading and reading-related skills: phonological processing (Comprehensive Test of Phonological Processing, CTOPP, Torgesen, Wagner, Rashotte, & Pearson, 1999), rapid naming (3-Set subtest of the RAN/RAS, Wolf & Denckla, 2005), single-word reading (Word ID and Word Attack subtests of the Woodcock Reading Mastery Test-Revised (WRMT-R, Woodcock, 2011), Passage Comprehension (WRMT-R, Woodcock, 2011), and the Reading Fluency subtest of the Woodcock-Johnson Test of Achievement Third Edition (WJ-III, Woodcock, McGrew, Mather, & Schrank, 2001). The performance on these assessments for all participants is summarized in **Table 1**.

**Table 1.** Mean (standard deviation) standard/scale scores for the reading and reading related subskills psychometric assessment.

|  | **1<sup>st</sup> time point** | **2<sup>nd</sup> time point** | ***t*** |
| --- | --- | --- | --- |
|  | Mean (SD) | Mean (SD) |  |
| Word ID | 109.85 (12.2) | 108.81 (10.73) | 0.73 |
| Word attack | 111.92 (15.81) | 106.6 (9.27) | 2.33 |
| Passage comprehension | 108.32 (12.4) | 109.23 (11.67) | -0.33 |
| Reading fluency | 105.48 (15.98) | 105.23 (13) | -0.04 |
| Phonological awareness- Elision | 11.04 (3.1) | 11 (2.93) | 0.07 |
| Phonological awareness-Blending | 10.65 (2.18) | 10.88 (2.88) | 0.57 |
| Phonological awareness- Nonword repetition | 9.81 (2.67) | 10.0 (2.97) | -0.33 |
| Rapid Alternating Stimulus Tests | 106 (19.27) | 104.19 (14.23) | 0.68 |

### Experimental tasks and imaging data analyses

#### Fluency task

This task was previously used and described by Benjamin & Gaab (2012) in adults and by Langer et al. (2013, 2019) in typically developing children and children with reading disabilities. For each trial, sentences comprised of four words were presented at a constrained, comfortable, or accelerated speed. The speed of word presentation in the constrained condition was fixed at 1350 ms for all participants. In contrast, the comfortable reading speed was customized for each participant outside the scanner (described in the subsequent section). The speed of the accelerated condition was 35% faster than the comfortable speed. As such, presentation speeds for comfortable and accelerated conditions varied across subjects and time points, while presentation speed for the constrained condition was the same across participants and time points. Word characteristics, including the age of acquisition, word frequency, familiarity, concreteness, imageability, and the number of phonemes and letters, were controlled using the MRC database (http://websites.psychology.uwa.edu.au/school/MRCDatabase/uwa_mrc.htm).

#### Determination of comfortable sentence reading speed

Before scanning, children underwent testing to determine their individual reading speeds. They were presented with three passages and asked to read them at a comfortable speed, taking as much time as necessary to complete. To capture their reading time, children pressed a key on a laptop to present each passage and another key when they finished reading the passage.

#### fMRI task

Before undergoing MRI, children underwent intensive training using a mock MRI scanner (for details, see Raschle et al., 2009, 2012). The fMRI implementation of the fluency task was identical to that used in Langer et al. (2013, 2019), a child-adapted version of the experimental fluency design employed in Benjamin & Gaab (2011). The task was presented in two nine-minute-long runs, which included real word sentence (i.e., task) and letter string sentence (i.e., control) conditions, each presented at constrained, comfortable, and accelerated speeds.

Participants were first presented with a picture cue indicating word presentation speed (turtle-constrained; cat-normal; rabbit-accelerated). Participants were then presented with a sentence one word at a time at one of the speeds (e.g., “The cat ran”), followed by a comprehension question. The comprehension phase included selecting one of three pictures that best describes the presented sentence. Children were instructed to choose the image that best represented the meaning of the sentence. For the control letter task, following the speed indicator picture, strings of ‘n’ letters were presented in place of the words, spaced to appear with a similar structure as sentences, with one different target letter. Children were asked to choose the oddball letter (“f,” “p,” or “x”) that appeared in one of the last two letter strings. This control task was designed to probe lower-level orthographic skills (e.g., visual attention/visual search) and letter recognition but not rely on high-level reading skills (e.g., semantic processing). Each of the two runs comprised 42 (21 words and 21 letter string) sentences, with the number of letters matched across conditions and runs. Across the two runs, 14 words and 14 letter string sentences appeared for each reading speed (constrained, comfortable, and accelerated).

Task and control trials were presented using an event-related design with the order of the two conditions (real word and letter string sentences) and speed pseudorandomized. Each trial began with an image cue indicating the upcoming presentation speed, which appeared on the screen for 500 ms and was followed by a black screen for 200 ms. Then, the words or control stimuli appeared from left to right at constrained, comfortable, or accelerated speed until the complete sentence was displayed. This was followed by a blank screen (200 ms). Subsequently, the comprehension or letter viewing testing phase appeared on the screen for 3000 ms or until the participant indicated their response (with a button press). The location of the correct image/letter was pseudorandomized in each trial. Each trial ended with a fixation cross presented for a variable time for up to 2000 ms. Performance was measured by the percent of trials answered correctly.

#### Imaging protocol and analysis

MRI scans were acquired on a SIEMENS 3.0T Trio MR whole-body scanner. 271 whole-brain images were acquired in each of the two fMRI runs with a 32-slice functional echo-planar acquisition (interleaved ascending) using TR=2000 ms, TE= 30 ms, FOV= 192 mm (full brain coverage), voxel size = 3 × 3 × 4 mm, and flip angle = 90°.

#### Preprocessing

The first four images of each run were discarded to account for field effects. Data were then preprocessed and analyzed using FSL 5.9 (http://www.fmrib.ox.ac.uk/fsl), beginning with motion correction (MCFLIRT), slice-timing correction, brain extraction (BET), linear registration (12 degrees of freedom) to the MNI 152 T1 template (FLIRT), spatial smoothing (4 mm FWHM kernel), and high-pass filtering (50 s). To deal with the relatively high degree of head motion common in pediatric neuroimaging, we used the ART toolbox (http://cibsr.stanford.edu/tools/human-brainproject/artrepair-software.html) to carefully detect volumes using a translation threshold of 2 mm and a rotation threshold of 0.02 mm. All subjects had two runs in which greater than or equal to 85% of the constituent volumes were free of artifactual volumes. Subjects not meeting this criterion were excluded from further analyses (*N* = 7). Motion parameters and artifactual volumes were entered as regressors in the first-level model.

#### fMRI analysis

Whole-brain analysis was performed in three stages. (1) A first-level model was designed for each participant and each run. Data were prewhitened and regressors were modeled for the speed cues; constrained, comfortable and accelerated fluent sentence reading; constrained, comfortable, and accelerated letter string reading; sentence and control comprehension stimuli; and intertrial fixation. Motion parameters and artifactual volumes were defined as confounding extraneous variables. (2) We used an event-related design in which the four words or letter strings constituted a single event. Note that unequal numbers of images were acquired for each participant and between the two time points since the individual reading speed varied between participants and between the two time points. FSL, however, can accommodate this variance (for a review, see Beckmann & Smith, 2004; Smith et al., 2004). Additionally, using FSL, low-level design matrices do not need to be identical to compare the subjects on a higher-level analysis ( Smith et al., 2004). (3) For each time point, the two-runs of each child’s data were combined in fixed-effects models and then entered into a group-level random-effects analysis (FLAME 1). Second-level statistical maps were generated using a (Gaussianized t-statistic) threshold of Z = 2.3 and a cluster-corrected threshold of P < 0.05 for the within-group and between-groups (i.e., time points) contrasts.

The following contrasts were examined (see also **Table 3**):

1. *Validation of sentence reading activation at time 1 and time 2.* To replicate previous results with children (Langer et al., 2015, 2019), we examined sentence reading activation at each speed (constrained, comfortable, accelerated) separately in contrast to fixation (rest condition) at both *time 1* and *time 2* reading stages. We then compared activation for each sentence speed condition (relative to fixation) between reading stages.
2. *Comparison between fluent sentence reading and letter string reading at time 1 and time 2*. We compared fluent sentence reading to letter string reading to identify brain regions that responded selectively to fluent sentence reading. We computed the contrast fluent sentence reading [all speeds] > letter string reading [all speeds], first for each reading stage separately, and then between the two reading stages (*time 2* > *time 1*).
3. *Comparison among reading speeds at time 1 and time 2 reading stages.* We compared activation at the two time points for conditions with higher presentation rate with conditions with lower presentation rate to identify brain regions that responded selectively to the increased demands of more rapid reading (sentence reading [accelerated] > sentence reading [comfortable]; sentence reading [comfortable] > sentence reading [constrained] and sentence reading [accelerated] > sentence reading [constrained]).

#### Region-of-Interest Analysis

Based on previous results using this paradigm (Benjamin and Gaab 2011; Langer 2013, 2019), a region-of-interest (ROI) analysis was performed for the bilateral vOT cortex. First, regions engaged in fluent sentence reading were identified through the contrast of sentence reading [comfortable] > sentence reading [constrained]. Second, ROIs were defined as the intersection between the functional activation and the fusiform region (one per hemisphere) as defined with the Harvard–Oxford anatomical atlas. Finally, subjects’ mean contrasts of parameter estimates (COPEs) were then extracted from ROIs for fluent sentence reading (comfortable > constrained) via *featquery* (http://www.FMRIb.ox.ac.uk/fsl/feat5/featquery.html) at each reading stage.

We then used these ROIs to investigate longitudinal brain-behavior associations. We first calculated the change in activation in the left and right vOT regions during fluent sentence reading (comfortable > constrained sentence reading) by subtracting the contrast maps of the *time 1* point from the *time 2* point for each participant. Next, we calculated the differences in raw scores between the two time points (*time 2*–*time 1*) for the WJ reading fluency test (Woodcock et al., 2001; a reading fluency measure) and the WRMT word attack subtest (Woodcock, 2011; a word decoding measure). Tests for partial correlations were performed between the brain and behavioral measures of reading controlling for differences in time passed between *time 1* and *time 2* behavioral and MRI data collection points, which varied across subjects.

## Results

### Psychometric assessment

Standardized psychometric test scores did not differ between *time 1* and *time 2* points (**Table 1**), according to a paired t-test, indicating that children retained their relative reading proficiency across time. However, as expected, raw psychometric scores differed between the two time points for all reading tests (Word ID, Word Attack, Passage Comprehension, and Reading Fluency) and the RAN (**Table 2**), indicating improved reading skills across time. No significant differences between 1^st^ and 2^nd^ time points were observed for phonological awareness as measured using the CTOPP raw scores.

**Table 2.**
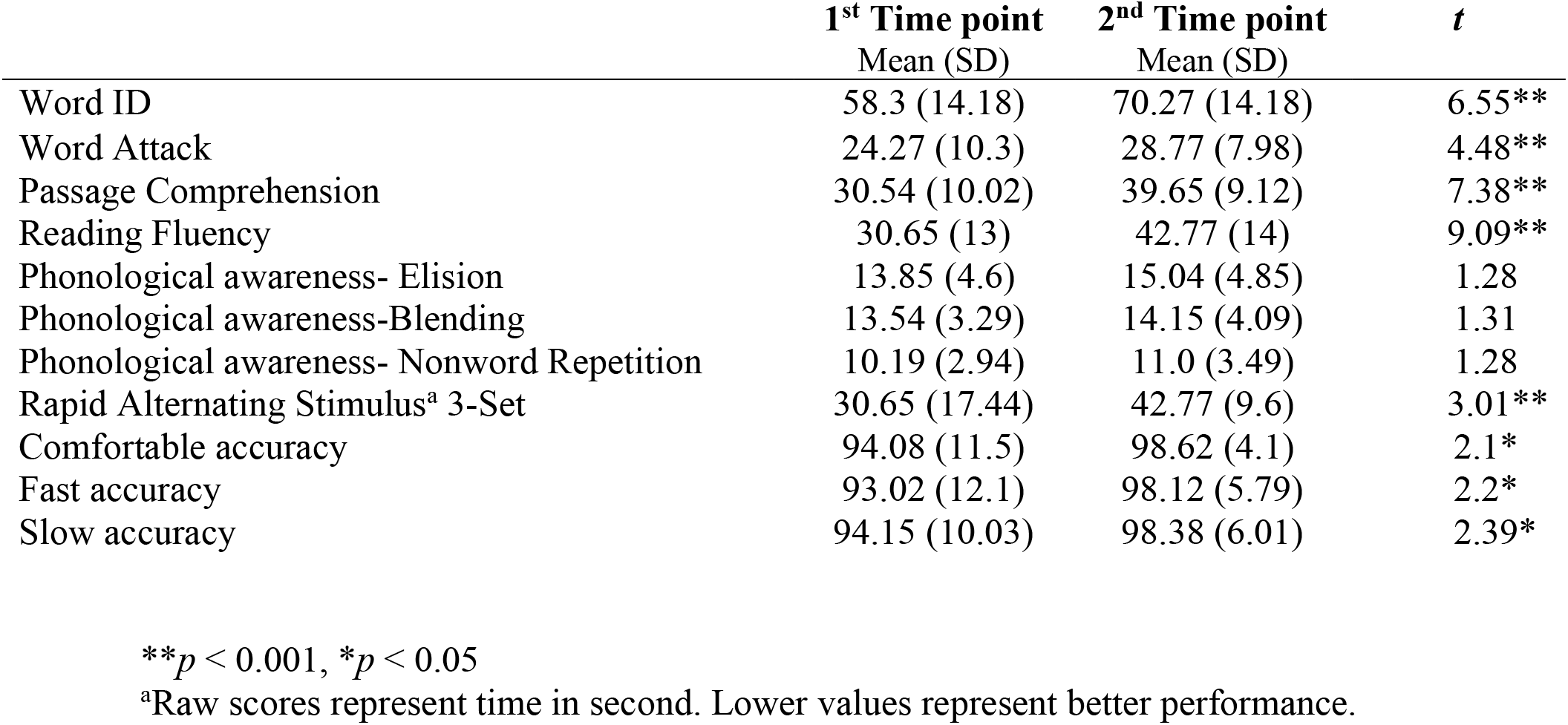
Raw scores (number of correct responses) for the psychometric assessments and in-scanner accuracy for each time point and t-score for the time points comparisons.

**Table 3.**
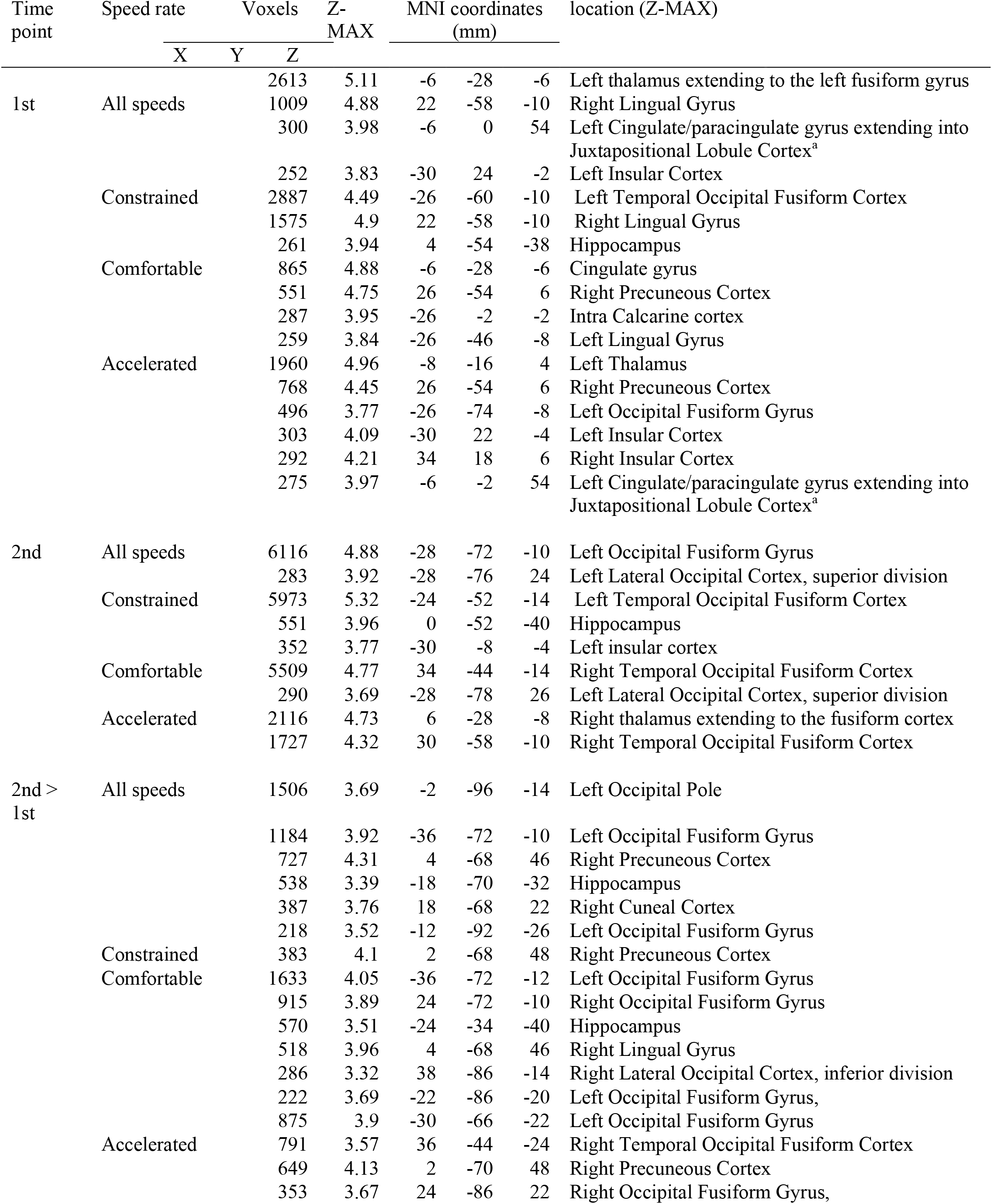
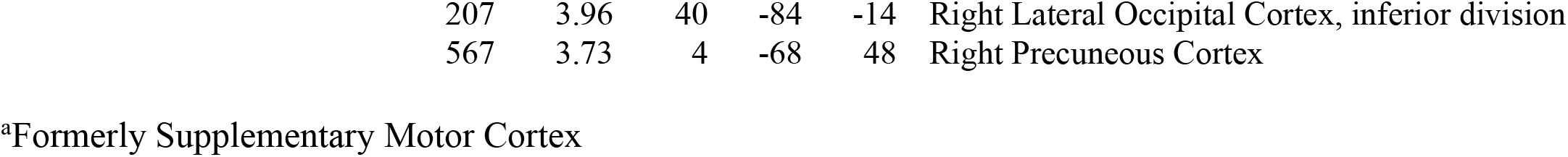
Results for the sentences reading (all speeds) > rest for each time point and the time points comparisons.

### Determining sentence reading speed

Comfortable reading speed, as determined before each MR scanning session, improved (i.e., reading rate increased) from *time 1* (ms/word = 621±288 ms) to *time 2* (ms/word = 454±132ms; *t*(25) = 4.03, *p* = 0.0001).

### In-scanner performance

We used a two-way repeated-measures ANOVA with time (*time 1* and *time 2*) and reading speed (constrained, comfortable, and accelerated) as within-subject variables to test for differences in sentence reading accuracy from *time 1* to *time 2* measurement. Results indicated a main effect of in-scanner reading accuracy [*F* (1, 25) = 6.244, *p* = 0.019] due to higher accuracy in the 2^nd^ time point (98±5%), compared to the 1^st^ time point (94±11%). Neither the effect of reading speed [*F* (2, 50) = 2.734, *p* = 0.075] nor the interaction between speed and time point *F* (2, 50) = 0.088, *p* = 0.918] were significant.

### fMRI results

#### Validation of sentence reading activation at time 1 and time 2

The results for these contrasts are presented in **Table 4**. When compared to rest, sentence reading (all speeds combined) for *time 1* and *time 2* activated the bilateral ventral occipito-temporal (vOT) regions, including the lingual gyrus and fusiform gyrus, and the insular cortex. A comparison between the two time points revealed increased activation at the *time 2* in the left fusiform gyrus for all three reading speeds (**Figure 1**).

**Figure 1.**
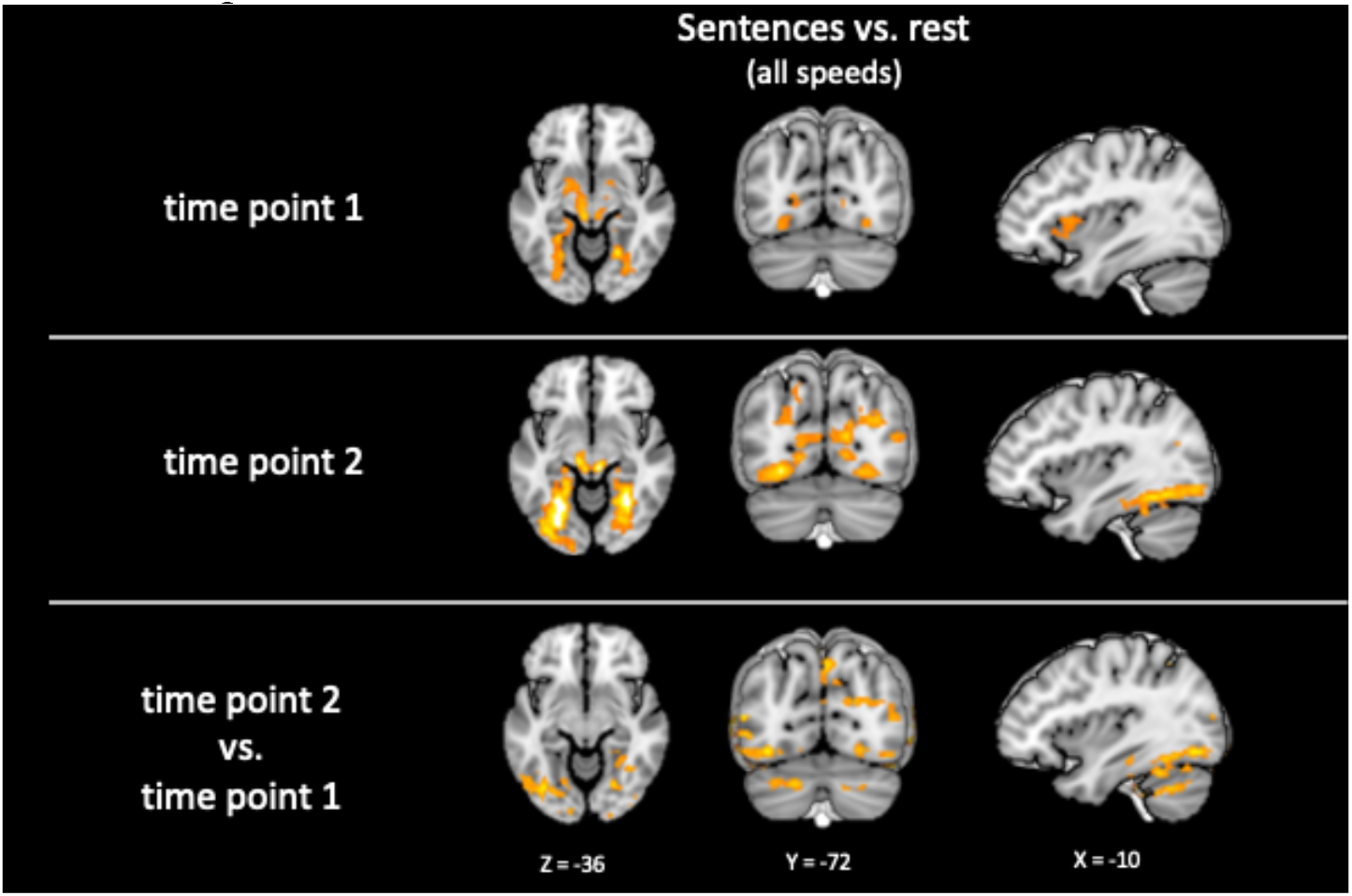
Fluent sentence reading (all speeds) > rest for the *time 1* point, *time 2* point, and the comparison between the two time points. Children show increased BOLD responses in several cortical and subcortical region mainly in occipito-temporal regions. The level of significance was set at *p* < 0.05 cluster-corrected.

**Table 4.**
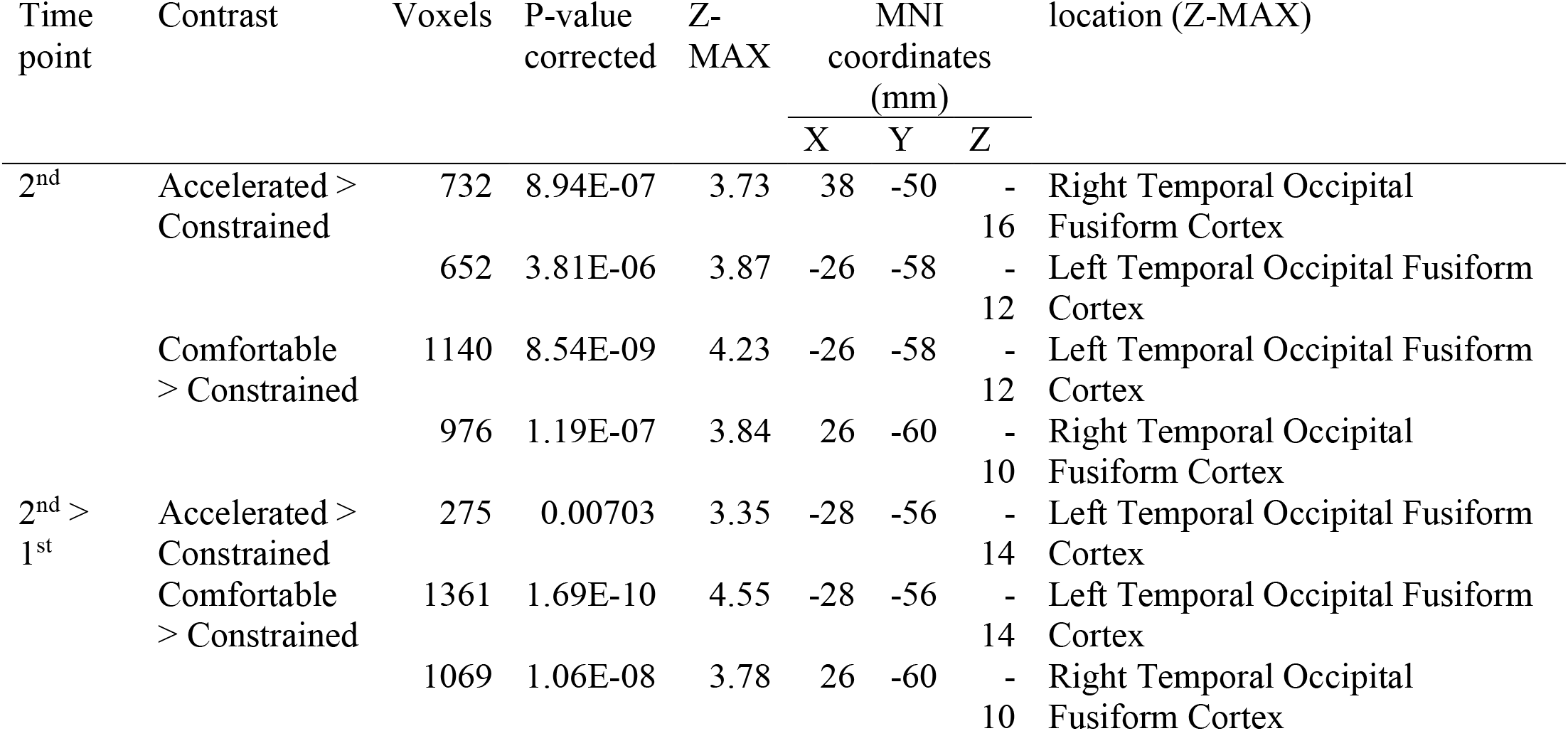
Results for the different sentences reading speed comparisons. No significant differences for the speed comparisons were found for the 1^st^ time point.

#### Comparison between fluent sentence reading and letter string reading at time 1 and time 2

The contrast of fluent sentence reading (all speeds) versus letter string reading (all speeds) revealed increased activation in the left and right fusiform cortex for both time points (**Figure 2**). A paired t-test for the contrast sentence > letters resulted in no significant differences between the two time points.

**Figure 2.**
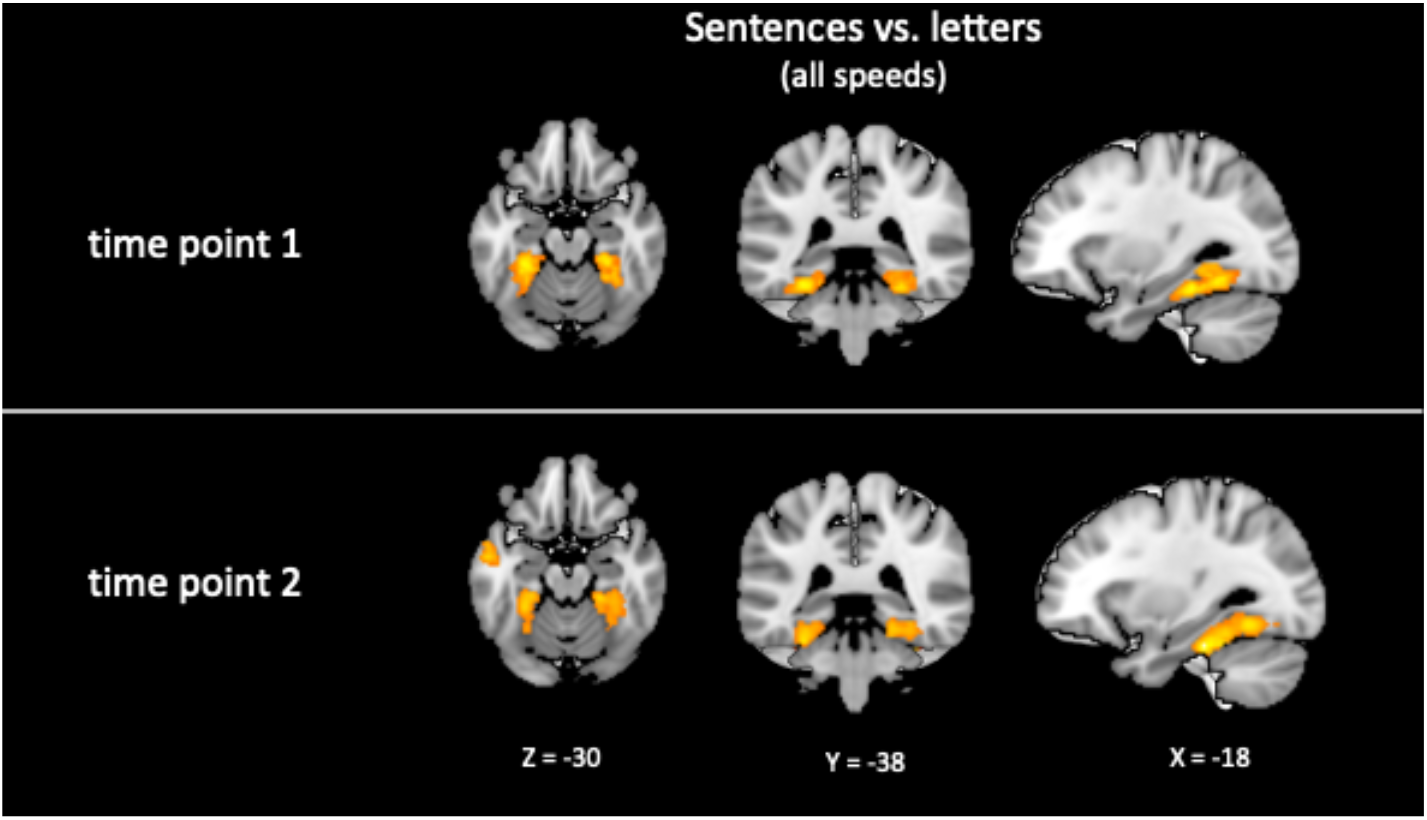
Fluent sentence reading (all speeds) > letter reading (all speeds) for *time 1* and *time 2*. Children show increased BOLD responses in bilateral ventral occipito-temporal (vOT) cortex for the sentence reading task vs. the letter reading task. Time points comparison did not reveal significant differences. The level of significance was set at *p* < 0.05 cluster-corrected.

#### Comparison among reading speeds at time 1 and time 2

Results for sentence reading rate contrasts (accelerated > constrained; comfortable > constrained; and accelerated > comfortable) are presented in **Table 5** and **Figure 3**. There were no significant differences in activation in response for the different contrasts for *time 1*. For *time 2*, greater activation was shown in the left fusiform cortex for accelerated > constrained contrasts and the bilateral fusiform cortex for comfortable > constrained contrast (**Figure 3**). No significant activations were found for the accelerated > comfortable contrast.

**Figure 3.**
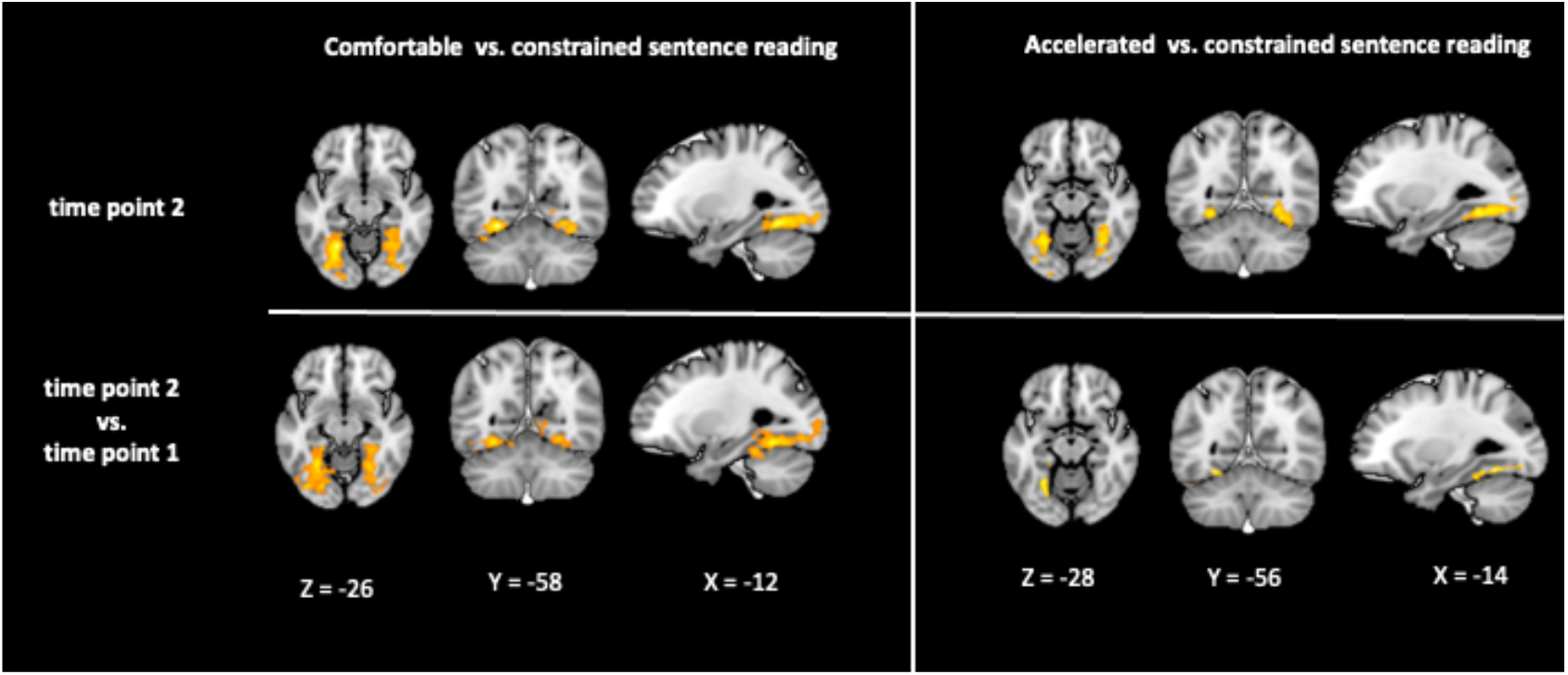
Comparisons of different reading speed (comfortable> constrained sentence reading; accelerated > constrained sentences reading) for *time 2* and the time points comparisons. Children show increased BOLD responses in bilateral vOT cortex in advanced reading stage and for the time points comparisons. No significant effects were found for *time 1*. The level of significance was set at *p* < 0.05 cluster-corrected.

For the longitudinal comparison, greater activation in *time 2* compared to *time 1* was shown in the bilateral fusiform cortex for the accelerated > constrained contrast and in the left fusiform cortex for the comfortable > constrained contrast.

#### Region of interest analysis

We tested six correlations between changes in activation for *time 2 – time 1* separately for left and right fusiform cortex and change in reading ability (raw scores for *time 2* - raw scores for *time 1*) while controlling for time passed between the two time points (please see Methods for details). A significant correlation (*r (26)* = 0.53, *p* < 0.05, Bonferroni corrected for multiple comparisons) was observed between change in left fusiform activation and change in reading fluency (measured by the WJ-III reading fluency subtest; **Figure 4**). No other relationships between brain function and reading abilities remained significant after correction for multiple comparisons.

**Figure 4.**
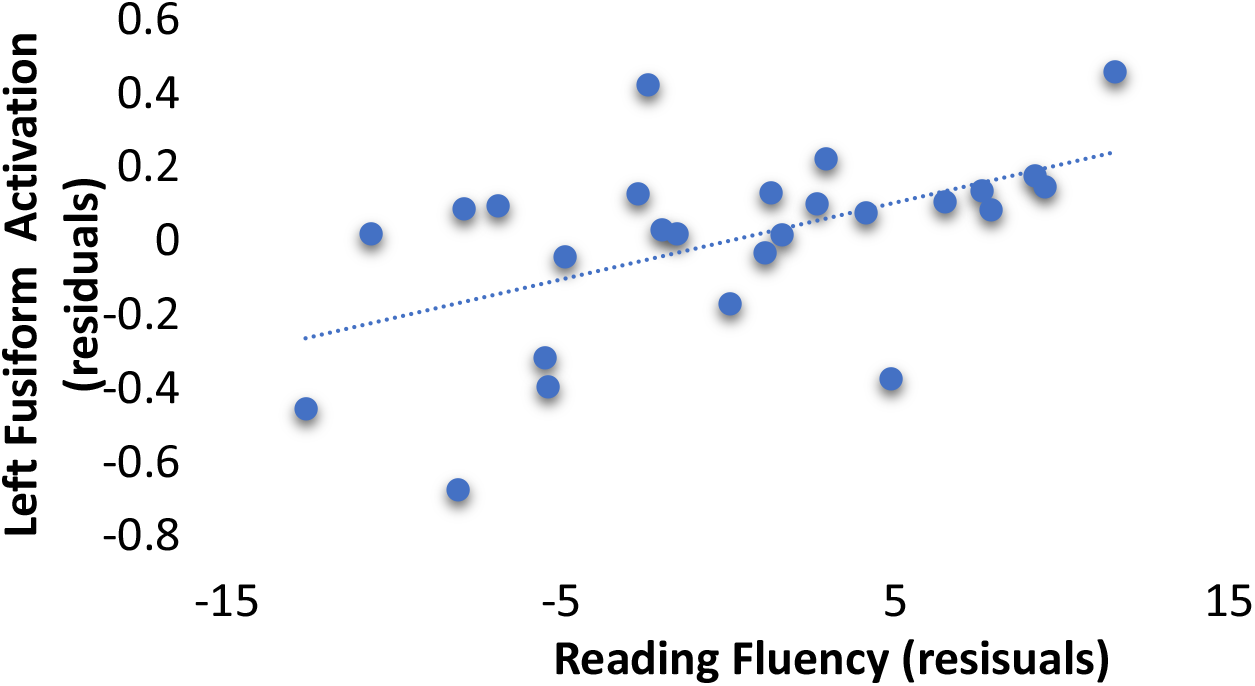
A scatter plot illustrating Partial correlation of growth in left vOT cortex activation (ROI) with growth in reading fluency (WJ-III) controlling for the time differences between two measures (*r (26)* = 0.53, *p* = .007).

**Figure 5.**
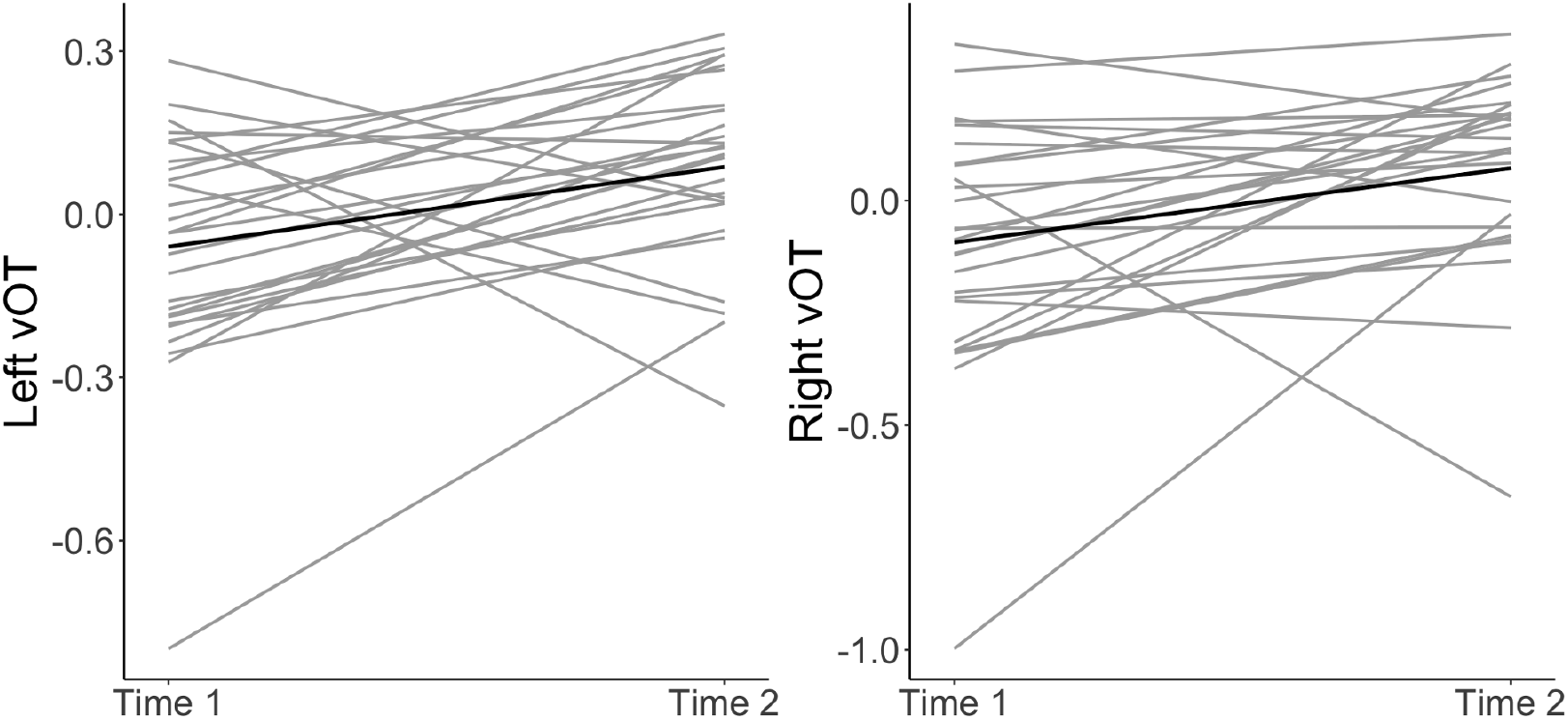
Individual growth lines for all participants representing increased activation from *time 1* to *time 2* in the left and right ventral occipito-temporal (vOT) regions.

## Discussion

The current study investigated developmental changes in neural patterns of activation underlying reading fluency. The study is novel in that it uses a longitudinal design and a more ecologically valid task that changes fluency demand by manipulating the presentation speed for each participant according to their individual-based reading speed. First, we demonstrated increased activation of the bilateral ventral occipito-temporal regions (vOT) in children during reading at their comfortable speed at *time 2* compared to *time 1.* Second, consistent with studies in same-age children (Christodoulou et al., 2014; Langer et al., 2013, 2019) and adults (Benjamin & Gaab, 2012), increased engagement of these regions was associated with increased speed demands at *time 2*, but not in *time 1*. Finally, increased activation in vOT was associated with a larger growth in reading fluency skills. Taken together, our findings provide critical insights on the association between the development of the vOT and children’s transition to fluent reading.

### Reading Activation Profiles at the Two Time Points

There were differences in activation patterns between time 1 and time 2 points, across all presentation speeds. Specifically, in *time 1*, children recruited insular, cingulate, and occipito-temporal regions during sentence reading. In *time 2*, there was significant recruitment of occipito-temporal areas only. The results in the older children were strikingly parallel to those obtained in previous studies using the same paradigm of comparable age or older individuals (Benjamin & Gaab, 2012; Langer et al., 2015, 2019). The younger group’s patterns of activation, however, were notably distinct (although the direct comparison did not reach significance), with increased recruitment of multi-demand regions during sentence reading (i.e., insular cortex, cingulate cortex, and precuneus). These regions were shown to support a range of executive control functions (e.g., inhibitory control, attentional selection, conflict resolution, maintenance and manipulation of task sets) for both linguistic and non-linguistic tasks (e.g., Duncan & Owen, 2000; Fedorenko et al., 2013; Hugdahl et al., 2015; see Fedorenko, 2014). Previous studies found increased engagement of these systems to support decoding in non-proficient readers (Roe et al., 2018; Ryherd et al., 2018; Ozernov-Palchik et al., 2020). These differences in patterns of activation between the two time points support the critical transition proposed around 3^rd^ grade from effortful reading that requires the utilization of considerable cognitive resources, to increasingly automatic word recognition. Such automatic word recognition is akin to the effortless processing of other visual objects such faces (Chall, 1996).

A direct comparison between the two time points revealed that activation in vOT increases longitudinally. Indeed, increased engagement of this region was shown to parallel increased perceptual expertise for processing words in longitudinal studies of early readers (Brem et al., 2010; Dehaene-Lambertz et al., 2018; Pleisch et al., 2019; Saygin et al., 2016). Specifically, as word processing in vOT becomes increasingly fluent, this region operates to rapidly extract invariant information from the word form, linking this information with the corresponding higher-level linguistic representations and attentional systems (Chen et al., 2019; Price and Devlin, 2011; Schlaggar and McCandliss 2007). Although previous studies demonstrated developmental differences in the region, this is the first study to adjust for potential differences in fluency-related task difficulty in a longitudinal design. This is important because of the sensitivity of vOT to differences in task demands and durations of exposure (Benjamin & Gaab, 2012; Dehaene & Cohen, 2011). By establishing each participant’s comfortable reading speed and choosing simple sentence stimuli, we equated task demands and the optimal exposure speed between the two time points, and across individuals. We can therefore confidently interpret our findings as representing increased specialization of the vOT region for fluent reading.

### Early Specialization of vOT for Reading

The contrast sentence > letter reading (averaged across all speeds) revealed similar patterns of activation in the bilateral vOT in both *times 1* and *2*. Our results support previous findings of print-induced activation in the vOT region at the beginning reading stages and its increase with reading experience (Centanni et al., 2017; Dehaene - Lambertz et al., 2018; Lochy et al., 2016; Saygin et al., 2016). As the region’s activation for single letters decreases, its activation for words increases. Specifically, a recent study has documented an inverted U-shaped pattern of activity of vOT to letter stimuli that peaks in the 1^st^ grade once the alphabetic code has been mastered, but then begins to dip (Fraga-Gonzales et al., 2021). Specialization for words follows a similar trajectory (Fraga González et al., 2014; Maurer et al., 2006). The onset of the vOT activation curve follows the decline in its responses to letters, and with a more extended peak, as word mastery is a longer milestone to reach (Centanni et al., 2017). Since previous developmental studies examined words presented in isolation, it remains unknown whether the course of specialization of vOT to connected text follows a similar trajectory. Our study precludes us from establishing the trajectory of the sentence-responses beyond our 2^nd^ time point (3^rd^/4^th^ grades). Longitudinal studies with additional measurement occasions are therefore needed to map out the course of vOT activation under naturalistic reading conditions.

The increased response in the current study of vOT to sentences, as compared to letters, is consistent with the Interactive Account (Price & Devlin, 2011) of vOT specialization. This account applies a predictive coding framework to describe how high-order language regions (e.g., phonological and semantic regions) generate predictions regarding the identity of words based on the contextual cues and lower-level visuospatial features. Activation in the vOT region reflects prediction error. i.e., the discrepancy between the predictive and the sensory signals. Predictions are stronger for words, especially when these words are embedded in sentences that provide strong contextual cues to word identification than for letter strings, resulting in more robust error signals and increased activation for words as compared to letters (Price et al., 1996).

This framework has important implications for developmental differences within the vOT. In preliterate children, vOT activation is low because the orthographic inputs fail to trigger the corresponding higher-level representations; therefore top-down influences are weak. In early readers, the discrepancy between the top-down and bottom-up signals is maximal, resulting in the strongest activation. The prediction errors will decrease with increased reading expertise. The exact timing of the activation peak in the vOT is difficult to ascertain precisely. Various confounding factors could affect activation patterns in the vOT. Transparency of orthography is one such factor (Aro & Wimmer, 2003; Carioti et al., 2021). For example, maximal activity was demonstrated in 2^nd^ grade in children reading in German, which has a more transparent orthography than English (Maurer et al., 2006; van der Mark et al., 2011). Another factor is processing demands imposed by the task. For example, if task demands are not controlled for, some of the vOT signal in younger readers could reflect additional effort. Additional factors include stimulus exposure durations (Schuster et al., 2015) and nature of the task (e.g., lexical decision, over or covert naming, silent reading, single words) that have varied across different studies and have implications for the generalizability of findings to natural reading (Rayner, 1998; Wehbe et al., 2014; Yarkoni, Speer, Balota, et al., 2008). It is therefore likely, based on the current findings, that the activation peak in children learning to read English orthography will coincide with the development reading framework that posits that reading mastery is achieved at around 4^th^ grade (Chall, 1996). Thus, increased activation in vOT in more expert readers, as compared to their younger counterparts, is consistent with the interactive specialization framework. This framework sees increased specialization of the region through recurrent connectivity between the linguistic and the vOT components (i.e., higher language and lower sensory levels) through the experience of learning to read.

How does fluency play into this framework? We showed that reading speed modulates brain activation only at the *time 2* point, after children became more proficient in reading. It is possible that increased speed demands resulted in more prediction errors due to less accurate top-down predictions. Since the predictive mechanisms are less developed in younger readers, faster presentation rates did not increase error signals in this group. Additionally, children in *time 1*, but not *time 2*, activated the cognitive control regions to a greater extent in the accelerated condition. As discussed above, this suggested increased recruitment of the multi-demand domain-general regions for executive control in response to higher task demands. Therefore, increased fluency demands increased processing within the vOT for the more proficient readers and increased cognitive control recruitment in the emerging readers.

### vOT Specialization in Relation to Fluency Skills

The changes in vOT activation across the two time points were related to improvements in reading fluency performance. These findings highlight the reciprocal interaction between reading development and the neural specialization for reading. As children become more proficient readers, vOT activation increases. As vOT becomes more specialized for reading, children become increasingly automatic in their word identification. The majority of the previous studies demonstrating links between reading proficiency and vOT specialization have focused on cross-sectional comparisons (e.g., Carreiras et al., 2009; Church, Coalson, Lugar, Petersen, & Schlaggar, 2008; Dehaene et al., 2010; Turkeltaub et al., 2003) or lower-level reading skills (e.g., Ben-Shachar, Dougherty, Deutsch, & Wandell, 2011; Maurer et al., 2006, 2011; Skeide et al., 2017). Our findings are important because they demonstrate that variable and individually determined increased fluency demands modulate the specialization of vOT for reading only after word identification proficiency is achieved.

According to Chall’s stages of reading, fluency is a bridge that moves students from proficient decoding to the extraction of meaning from connected text (Chall, 1983; Chard et al., 2002). The transition from a focus on accurate word identification to using text to gain new knowledge and ideas through reading increasingly complex texts (reading to learn) is set to occur in 4^th^ grade in English-readers. Our findings that beyond just proficiency in decoding, rate of decoding is an important contributor to vOT activity, supports the significant role of fluency in reading development. We show that activation of vOT is a sensitive index of speed of processing -- and consequently of automaticity in word recognition -- that underlies the development of fluency in this critical period of transitioning into reading fluency and proficiency. In accordance with the multi-componential view of fluency, vOT is an important hub that receives and processes predictive signals from higher-language brain regions that support semantic, phonological, and attentional processes as well as feed-forward lower-level visual signals. Our findings suggest that the automaticity of processing in this hub underlies fluent reading and that the transition to the “reading to learn” stage is a critical time for this region’s neural specialization.

The educational implications of individual variability in reading fluency are great. In one study of students who took the NAEP reading assessment in 2002, 40% of the fourth-grade sample were identified as “non-fluent” readers (Daane et al., 2005). Fluency skills account for a significant and unique variance in reading comprehension (Cutting et al., 2009; Joshi & Aaron, 2000; Silverman et al., 2012; Silverman et al., 2013; Tilstra et al., 2009), and mediates the relationship between word reading and reading comprehension (Kim, Quinn, & Petscher, 2021; Kim & Wagner, 2015). The negative impact of fluency deficits have been shown to extend to many other school subjects (NRP, 2000). Indeed, dysfluent reading diverts cognitive resources such as attention and working memory away from comprehension and thereby hinders deep processing of and learning from text (Cain et al., 2004; Perfetti, 1985). Furthermore, reading speed is strongly associated with children’s self-concept regarding their reading skills and affects their motivation for reading (Kasperski et al., 2016).

Our findings can be extended to support the significance of interventions that prioritize fluency-building strategies. Repeated reading is the most common strategy that explicitly targets reading fluency. In a recent meta-analysis 90% of fluency intervention studies focused on this strategy (Hudson et al., 2020). These interventions aim to both promote fluency and advance more distal outcomes of reading comprehension. Based on the neurocognitive development model of vOT specialization, repeated reading would strengthen the connections between visual word features and their higher-level linguistic counterparts, resulting in increased automaticity of processing of orthographic patterns and subsequently greater fluency. Greater fluency would allow the multi-demand network to be engaged in the cognitively demanding process of assigning meaning, monitoring, inferring, and building coherence while reading connected text.

## Conclusions

We examined the development of the neural correlates underlying the development of reading fluency throughout 1-2 years of elementary schooling in a longitudinal design. Our results showed increased ventral occipito-temporal activation longitudinally and when increasing reading speed demands. Furthermore, increased activation was associated with better fluency development. These findings shed light on the reciprocal importance of the ventral occipito-temporal cortex for the development of reading fluency. Specifically, the increased engagement of this region in sentence reading (compared with letter strings) and during accelerated reading is modulated by and supports reading proficiency. These findings also provide mechanistic insights for the efficacy of repeated reading strategies to increase reading fluency.

## Acknowledgments

This work was supported by a grant from the National Institutes of Health–National Institute of Child Health and Human Development (Grants R01HD067312 to NG, F32-HD100064 to OO). This work was also supported by the Jacobs foundation and Charles Hood foundation grants to NG, and Harvard Brain Initiative Transitions Program grant to TKT. We sincerely thank our research testers and participating families. The authors have no conflicts of interest to report.

## Data Availability

The data required to reproduce reported findings will be provided upon request.

---

**Supplemental Table 1.**
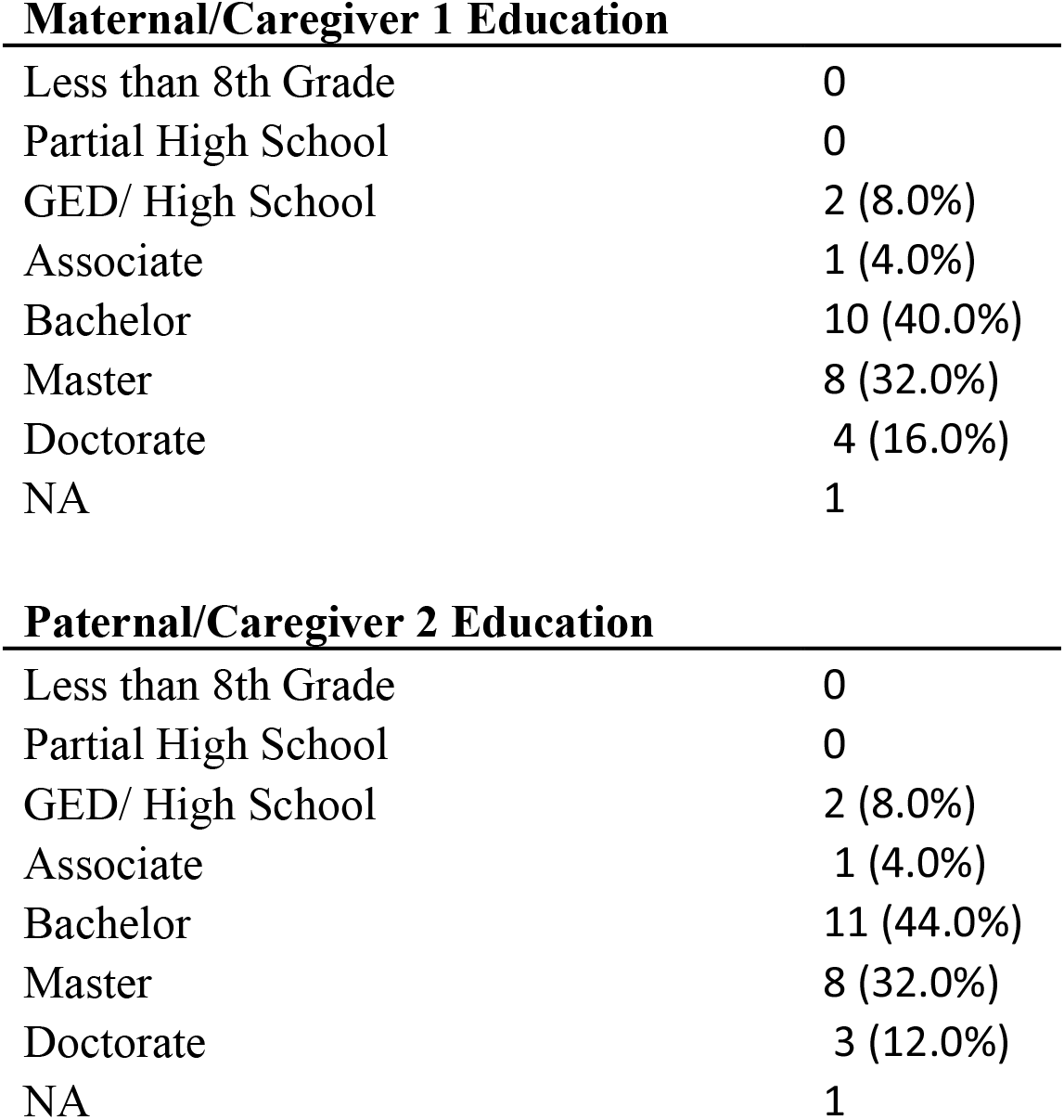
Participant Demographics (N=26)

**Maternal/Caregiver 1 Education**
|  |  |
| --- | --- |
| Less than 8th Grade | 0 |
| Partial High School | 0 |
| GED/ High School | 2 (8.0%) |
| Associate | 1 (4.0%) |
| Bachelor | 10 (40.0%) |
| Master | 8 (32.0%) |
| Doctorate | 4 (16.0%) |
| NA | 1 |
| **Paternal/Caregiver 2 Education** |  |
| Less than 8th Grade | 0 |
| Partial High School | 0 |
| GED/ High School | 2 (8.0%) |
| Associate | 1 (4.0%) |
| Bachelor | 11 (44.0%) |
| Master | 8 (32.0%) |
| Doctorate | 3 (12.0%) |
| NA | 1 |

## Notes

### Competing Interest Statement

The authors have declared no competing interest.

